# Optimal Strategies for Signal Sending and Perception in Volatile-mediated Within-Plant Signaling against Herbivory

**DOI:** 10.64898/2026.05.06.723397

**Authors:** Shuichi N. Kudo, Koki Iwakura, Akiko Satake

**Author notes:** Corresponding author: S. N. Kudo.

## Abstract

Herbivore-induced plant volatiles (HIPVs) play a critical role in inducible plant defense as information-bearing airborne signals. Released from damaged tissues, HIPVs induce defense responses in undamaged parts of the plant, thereby reducing the risk of subsequent herbivore attack. Although both emission and perception are fundamental components of HIPV-mediated signaling, the co-evolutionary dynamics of these traits under herbivore-driven selection remain poorly understood. Here, we develop a mathematical model of within-plant signaling that explicitly incorporates both inducible signal emission and perception as evolving traits. Using the model, we derived the optimal level of HIPV signal emission and signal perception under successive herbivore attacks. Our results show that the strategy with both signal emission and signal perception, which underlies HIPV-mediated signaling, is favored only under intermediate levels of herbivory. Within this range, increasing herbivory frequency drives the joint evolution of reduced signal emission and enhanced sensitivity to released signal. Furthermore, extending the model to include perception-independent functions of HIPVs, such as the attraction of natural enemies and the deterrence of herbivores, expands the range of conditions under which HIPV-mediated signaling is favored. At the same time, it also allows the emergence of emission-only strategies lacking signal perception, suggesting the potential decoupling of the co-evolution of emission and responsiveness. These findings provide a theoretical framework for understanding how emission and perception jointly shape the evolution of volatile-mediated signaling systems in plants.

## 1. Introduction

Herbivore-induced plant volatiles (HIPVs) are an important component of inducible defense mechanisms in plants (Frost et al., 2008; Heil & Karban, 2010; Heil & Silva Bueno, 2007). They comprise a diverse array of secondary metabolites, including monoterpenes, sesquiterpenes, and green leaf volatiles (GLVs) (Dudareva et al., 2004). Some HIPVs are produced prior to damage and stored in specialized structures such as oil glands and resin ducts, from which they are released upon mechanical damage caused by herbivores (Niinemets, 2018), while others are synthesized de novo in response to insect feeding (War et al., 2011). HIPVs function as information-bearing chemical signals: once released from damaged tissues, they are transmitted through the air and induce defense responses in other parts of the same plant, reducing the risk of subsequent herbivore attack (Frost et al., 2007; Karban et al., 2006). Such airborne signaling enables rapid information transfer even between branches that are not directly connected by vascular tissues, compared to vascular signal transduction (Frost et al., 2008). In addition to this within-plant airborne signaling among branches, HIPVs are also perceived by neighboring plants (plant–plant communication), herbivores, and their predators (carnivore–plant communication). In particular, they serve as cues for predators that indicate the presence of herbivores, thereby attracting natural enemies as bodyguards (Dicke & Baldwin, 2010; Takabayashi, 2022; Takabayashi & Dicke, 1996).

There is a substantial intra- and interspecific variation in HIPVs emission (Dicke & Baldwin, 2010). For instance, the total amount of HIPVs emission has been reported to vary up to eightfold among maize cultivars, as well as among their wild ancestors, teosinte (Gouinguené et al., 2001). Such quantitative variation has also been documented across various plant species including cotton (Loughrin et al., 1995), *Datura* (Hare, 2007), and *Nicotiana* species (Schuman et al., 2009). Because HIPVs mediates defense response against herbivores, differences in selective pressure by herbivores has been proposed as a key driver of evolution of HIPVs emission (Dicke & Baldwin, 2010). Consistent with this idea, invasive populations of ragwort, which have been released from selective pressure by specialist herbivores, exhibited reduced HIPV emission compared with populations in their native range (Lin et al., 2021). However, contrasting evidence has also been reported: a meta-analysis of 236 experiments from 109 studies showed that cultivated plants—presumably subjected to reduced herbivory due to insecticide use— emit higher levels of HIPVs than wild plants (Rowen & Kaplan, 2016). These findings highlight the complex relationship between herbivore pressure and the evolution of HIPV emissions.

In contrast to emission traits, variation in HIPVs perception traits, which determine how sensitively plants respond to HIPVs, remain largely unexplored. In animals, the coevolution of signal production and perception traits has been explored using several signaling systems such as insect pheromones (Groot et al., 2016; Symonds & Elgar, 2008) and mammal vocal communications (Charlton et al., 2019). For example, diversification of female sex pheromones in moths has been associated with the evolution of male preference (Boake, 1991; Groot et al., 2016). In acoustic communication among forest mammals, vocal characteristics and hearing sensitivity have co-evolved to utilize higher frequency sound, which are effective for localizing signals to avoid the detection by predators in visually occluded environments (Charlton et al., 2019). These findings from animal communication research indicate the HIPVs emission traits and perception traits may likewise undergo coevolution in plants. However, due to the difficulty in directly measuring perception traits and limited understanding of the molecular basis of HIPVs perception, how herbivory drives the coevolution of HIPVs emission and perception is still unclear.

To elucidate the driving factors and conditions underlying the evolution of HIPVs emission strategies, mathematical modeling approaches have been applied, primally focusing on tritrophic interaction-in which plants attract natural enemies of herbivores (Kobayashi et al., 2006; Kobayashi & Yamamura, 2003; Yamauchi et al., 2015)- and plant-plant communication (Hirose & Satake, 2024). These studies, grounded in evolutionary game theory, have provided theoretical insight into HIPV emission strategies from the perspectives of cooperative behavior (Kobayashi & Yamamura, 2003), signaling theory (Yamauchi et al., 2015), and public goods game (Hirose & Satake, 2024). Other frameworks have drawn on optimal defense theory (Fagerström et al., 1987; Iwasa et al., 2025; McKey, 1974). For example, it has been demonstrated that behavioral characteristics of herbivores influence the evolution of induced defense strategy mediated by volatile signaling and vascular transport based on the trade-offs between the costs and benefits of defense investments (Ito & Sakai, 2009). However, in these models, responsiveness to HIPVs is implicitly treated as a fixed, already evolved trait. Therefore, it is essential to advance research that explicitly incorporates HIPV perception as an evolving trait, thereby enabling a deeper understanding of the coevolutionary dynamics between emission and perception traits that shape both within-plant signaling and plant-plant communication mediated by HIPVs. Here, we develop a mathematical model of within-plant signaling that explicitly incorporates both induced signal emission (HIPV emission in response to herbivory) and perception (HIPV-dependent defense induction) under successive herbivore attacks. Using this model, we derive the optimal amount of HIPVs emission and the optimal response threshold for HIPV-dependent defense induction by minimizing total biomass loss, which accounts for both herbivore damage and the costs associated with signal production and defense induction. Our results show that strategies combining induced signal emission and signal perception are optimal only under intermediate levels of herbivory. Within this range, increasing herbivory frequency leads to the joint evolution of reduced emission and lower response thresholds, resulting in heightened sensitivity to HIPVs. Furthermore, incorporating perception-independent benefits, such as the attraction of natural enemies and direct deterrence of herbivores, expands the conditions favoring HIPV-mediated signaling, while it also promotes the emergence of emission-only strategies, suggesting a potential decoupling of the evolution of emission and responsiveness.

## 2. Model

### 2.1. Modeling emission and perception of volatiles in response to herbivory

To formulate volatile-mediated signaling within an individual plant, we consider two key processes: (i) HIPV emission from a damaged branch and (ii) volatile perception, whereby HIPVs induce defense responses in undamaged branches (Fig. 1). When a branch experiences herbivory, it emits an amount *e* ≥ 0 of volatiles to signal other branches within the same individual. Then, we define the state of the plant as *x* = 1 when it is experiencing herbivory and *x* = 0 otherwise, and the volatile emission from the branch can be expressed as

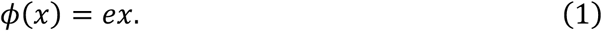

**Figure 1.**
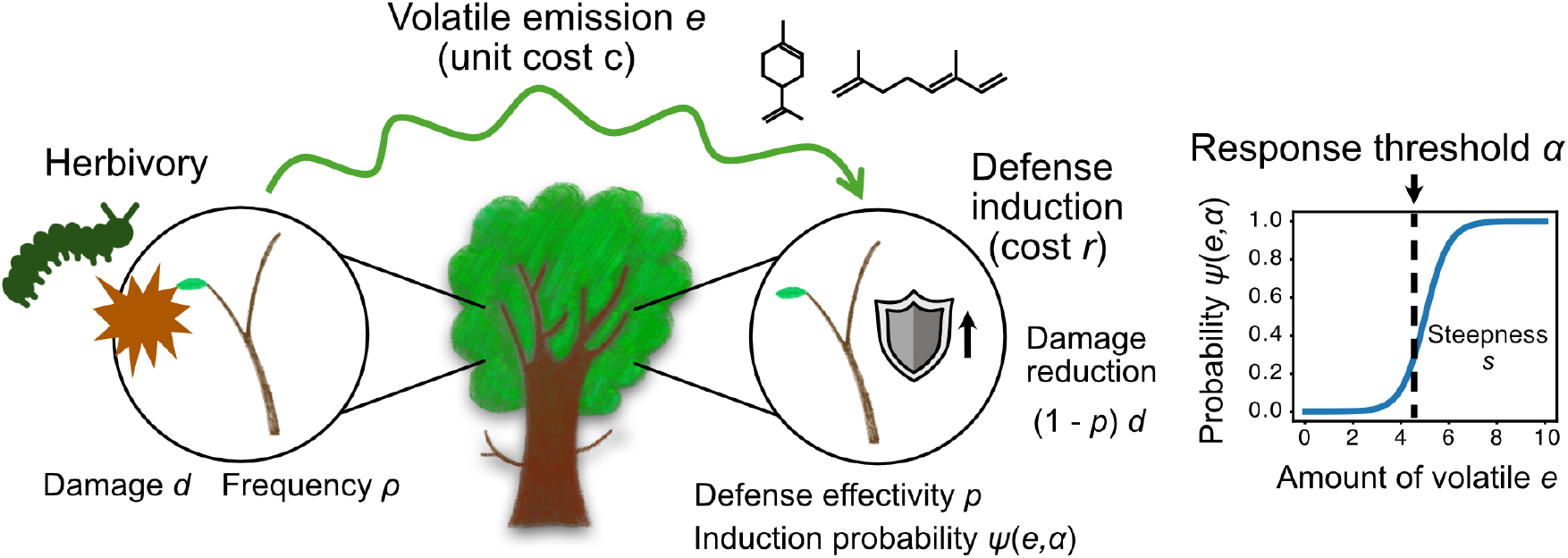
Schematic representation of the mathematical model of volatile-mediated within-plant signaling. Damaged branches emit volatiles at rate *e* and the other branches induce defense responses with a probability *ψ* determined by the emission *e* and the response threshold *α*.

In this paper, we consider the simplest case in which the plant emits a constant amount of volatile *e* only when it is damaged by herbivory.

To model volatile perception process and subsequent defense induction, we describe the probability of triggering a defense response in an undamaged branch as a function of the received volatile signal *φ*, bounded between 0 and 1:

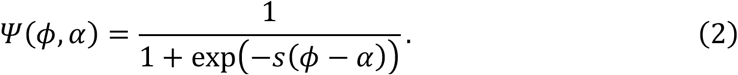

We assume a sigmoidal response function, such that the probability of defense induction increases with the amount of volatile received. The parameter *s* ≥ 0 determines the steepness of the response curve, and − ∞ ≤ *α* ≤ ∞ represent the response threshold. Biologically, *α* characterizes the sensitivity of volatile perception: lower values of *α* correspond to higher responsiveness to volatile cues. In the limiting cases, extreme values of *α* correspond to scenarios in which the probability of defense induction becomes independent of HIPV signals. As *α*→∞, the probability of defense induction *ψ* approaches zero regardless of volatile concentration, and plants do not induce defense even in the presence of HIPVs (“silent”). Conversely, as *α*→−∞, *ψ* approaches one regardless of the amount of HIPVs released, and plants constitutively express defense responses even in the absence of HIPVs (“constitutive defense”). Importantly, we treat the emission level *e* and the respond threshold *α* as evolutionary traits governing signal production and signal perception, respectively.

### 2.2. Definition of cost function based on expected carbon loss

We assumed that the focal individual is subjected to herbivory with a probability 0 ≤ *ρ* ≤ 1 at each time (Fig. 1). When the plant is attacked by herbivores, it loses an amount of biomass *d* ≥ 0. This damage is reduced by a proportion of 0 ≤ *p* ≤ 1 if the defense response is induced. Here we define the state of defense induction as *y* = 1 when defense is induced with the probability given by equation (2) and *y* = 0 otherwise. Using the state function of *x* and *y*, we can define the loss of biomass due to herbivory as:

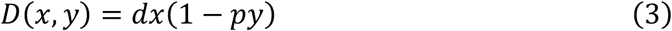

Next, we consider the costs associated with signal production *C*_*s*,_ and defense *C*_*d*_ as follows:

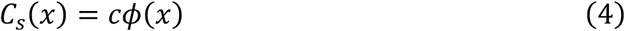

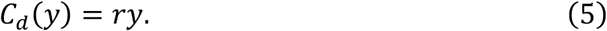

Because HIPVs are derived from photoassimilates, higher levels of emission require greater carbon and energy investment, resulting in metabolic costs (Sharkey et al., 2008). In addition, increased HIPV emission may enhance plant apparency to herbivores, thereby elevating the risk of attack and imposing ecological costs (Kessler, 2015; Strauss et al., 2002). Accordingly, we assume that the cost of signal production is proportional to the level of HIPV emission.

Finally, we define the total biomass loss *L* as the expected value of the sum of herbivore damages and the cost for volatile emissions and defense responses for fixed parameters *ρ, d, p, c*, and *r* ≥ 0:

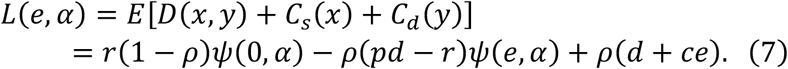

## 3. Result

### 3.1. Classification of the optimal strategies under different herbivory pressure

To derive the optimal volatile emission and response threshold for within-plant signaling, we first analyzed how the loss function *L* depends on the evolutionary traits *e* and *α*. Figure 2 illustrates contour plots of the loss function *L*(*e, α*), revealing three qualitatively distinct cases. Figure 2A represents a case in which a combination of a positive volatile emission rate and a finite response threshold minimize the carbon loss, corresponding to a *signaling strategy* (*e*_min_ > 0 and *α*_min_ >0). In contrast, Figure 2B depicts a case in which neither volatile emission nor defense response is favored, representing a *silent strategy* (*e*_min_ = 0 and *α*_min_ = ∞). Figure 2C corresponds to a *constitutive defense strategy*, in which plants always express defense responses regardless of volatile exposure (*e*_min_ = 0 and *α*_min_ = −∞). In the following sections, we demonstrate that these three strategies constitute the only possible global optima of the model. We further derive the conditions under which each strategy is favored and analytically determine the corresponding optimal volatile emission *e* and response threshold *α*.

**Figure 2.**
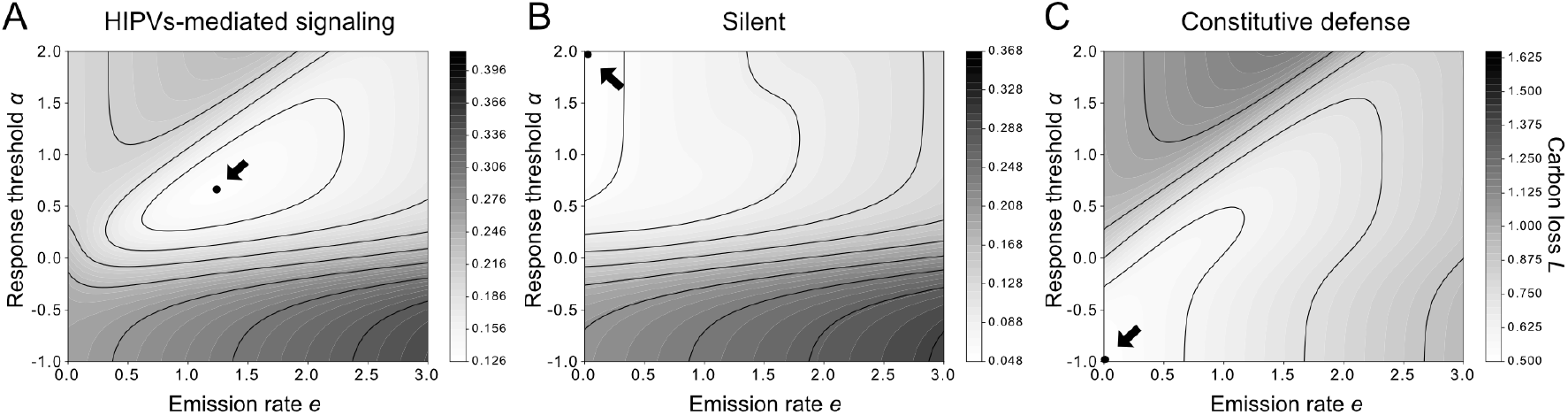
Contour plots of carbon loss function *L*(*e, α*). Black dots and arrows represent the minimum point of the loss. (A) Signaling strategy (*e*_min_ > 0 and *α*_min_ >0) is the optimum (*ρ* = 0.1, *d* = 2, *p* = 0.7, *r* = 0.2, *c* = 0.3, *s* = 5). (B) Silent strategy (*e*_min_ = 0 and *α*_min_ = ∞) is the optimum (*ρ* = 0.1, *d* = 0.5, *p* = 0.7, *r* = 0.2, *c* = 0.3, *s* = 5). (C) Constitutive defense strategy (*e*_min_ = 0 and *α*_min_ = −∞) is the optimum (*ρ* = 0.5, *d* = 2, *p* = 0.7, *r* = 0.2, *c* = 0.3, *s* = 5).

### 3.2. Optimal volatile emission and response threshold for defense induction

To derive the optimal signaling strategy of the focal plant individual, we obtain the minimum point of the cost function *L*. To this end, we compute the partial derivatives of *L* with respect to evolutionary traits, emission rate *e* and response threshold *α*, and determine (*e, α*) at which both derivatives simultaneously vanish. The partial derivatives of *L* are given as follows:

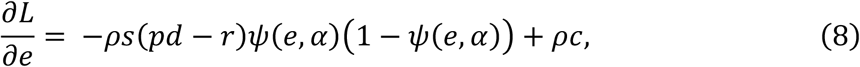

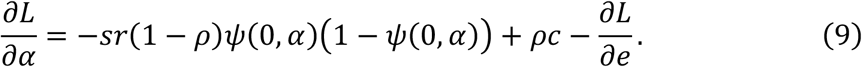

We denote (*e*^∗^, *α*^∗^) the value at which the partial derivatives vanish. Solving the equations for ψ(*e*^∗^, *α*^∗^)and ψ(0, *α*^∗^)yields

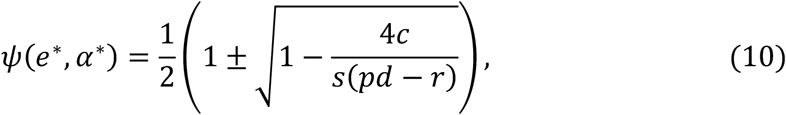

And

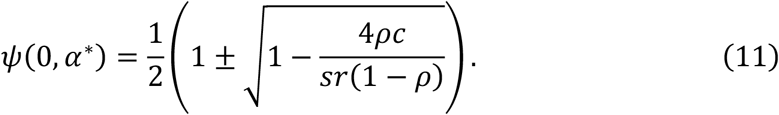

Moreover, the Hessian matrix of ψ at (*e*^∗^, *α*^∗^) is

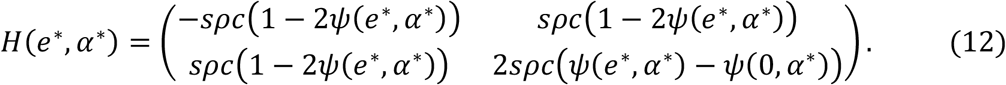

Hence,

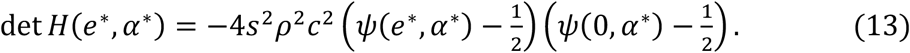

If (*e*^∗^, *α*^∗^)is a local minimal point of ψ, then *det H*(*e*^∗^, *α*^∗^)< 0. Together with the inequality ψ(*e*, α)> ψ(0, α)for all *e* and α, the values ψ(*e*^∗^, *α*^∗^)and ψ(0, *α*^∗^)are uniquely determined to be one of the two candidates,eq]

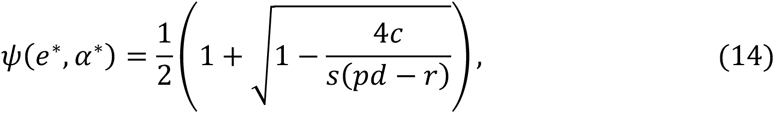

And

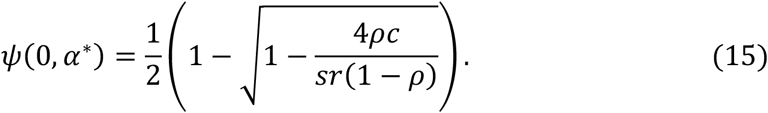

Here, *ψ*(*e*^∗^, *α*^∗^) represent the optimal defense induction probability in response to volatile exposure and *ψ*(0, *α*^∗^) indicates the basal defense probability without volatiles, respectively. The solutions (14) and (15) exist only when the terms in the square roots are positive. Therefore, the conditions for their existence are given by the following two inequalities:

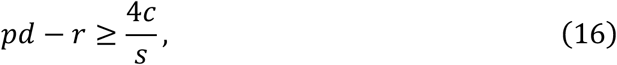

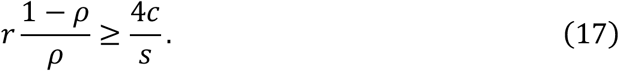

The left-hand side of inequality (16) represents the relative benefit of defense, reflecting the balance between the benefit of damage avoidance, *pd*, and the cost of defense induction, *r*, triggered by volatile signals. In contrast, inequality (17) characterizes the advantage of a conditional defense response mediated by volatiles. Because the term *r*(1−*ρ*) corresponds to the cost of inducing defense in the absence of herbivory, a large value of *r*(1−*ρ*) makes it advantageous to suppress defense induction when herbivory does not occur. This, in turn, favors the use of volatile-mediated signaling to achieve conditional defense responses specifically in response to herbivore attacks.

Hereafter, we denote *X* and *Y* as

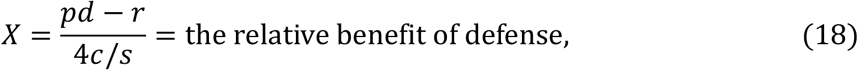

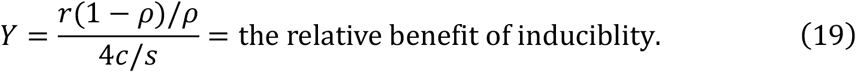

Inequalities (16) and (17) can be rewritten as *X* ≥1 and *Y* ≥1. Finally, based on the Eq. (2), we obtain the local minimum (*e*^∗^, *α*^∗^) of *L* using the inverse of the sigmoidal function,

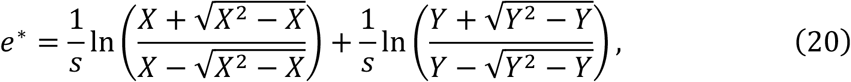

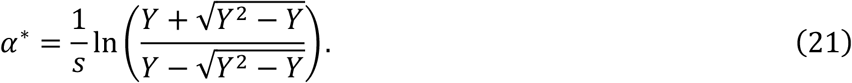

If inequality (16) or (17) is not satisfied, the carbon loss *L* becomes minimum on the boundary *e* = 0. By substituting *e* = 0 into *L*, we get

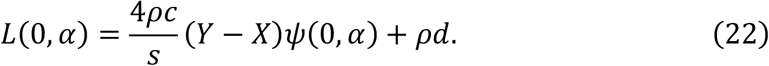

The equation above shows that cost *L* is a linear function of basal defense probability *ψ*(0, *α*). Thus, the optimal value *α* depends on the sign of the slope, which is determined by the balance between the relative benefit *X* and *Y*. Then, the value is as

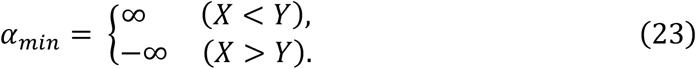

Note that, in case of *X*=*Y, L* is independent of *α*. The above result indicates that when herbivory occurs sufficiently infrequently (large Y), and the relative benefit of defense induction is small (small *X*), the silent strategy, (*e,α*) = (0,∞) —characterized by the absence of volatile emission and defense induction—becomes optimal (Fig. 2B). In contrast, under frequent herbivory (small *Y*) and a large benefit of defense induction (large *X*), the constitutive defense strategy, (*e,α*) = (0,−∞), in which plants continuously induce defense responses without relying on volatile signals, is favored (Fig. 2C).

Figure 3 illustrate how the optimal HIPVs emission *e*^∗^ and response threshold *α*^∗^ depend on the leaf damage *d*, which increases relative benefit of defense *X*, and on herbivory frequency *ρ*, which decreases the relative benefit of inducibility *Y*. First, increasing damage severity *d* promotes higher HIPVs emission while the optimal response threshold remains unchanged (Fig. 3A). Consequently, the separation between defense expression under herbivory *ψ*(*e*^∗^, *α*^∗^) and in its absence *ψ*(0, *α*^∗^) becomes larger, indicating enhanced inducibility (Fig. 3B). In contrast, herbivory frequency *ρ* affects both emission *e*^∗^ and perception traits *α*^∗^ (Fig. 3C). As *ρ* increases, the optimal emission level *e*^∗^ and the response threshold *α*^∗^ decrease simultaneously (Fig. 3C). This coordinated reduction maintains inducible defense *ψ*(*e*^∗^, *α*^∗^) while lowering the overall costs of signaling and defense induction under more frequent attack (Fig. 3D). When herbivory becomes sufficiently frequent, the optimal emission level eventually declines to zero (Fig. 3C), and the plant shifts to constitutive defense *ψ*(*e*^∗^, *α*^∗^) = *ψ*(0, −∞) = 1, expressing defense responses independently of volatile signaling (Fig. 3D).

**Figure 3.**
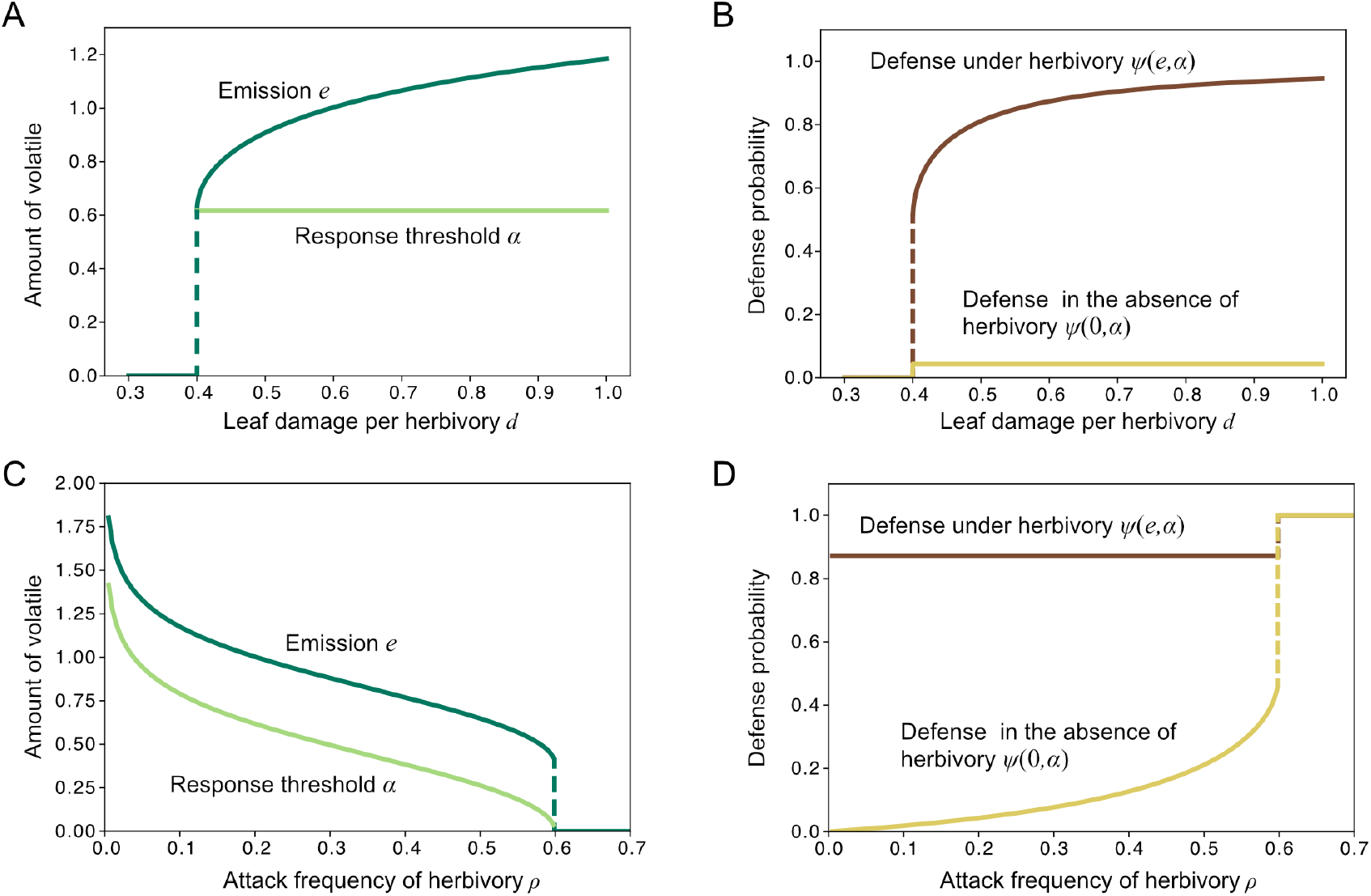
Parameter dependency of the optimal HIPVs emission and response threshold. (A) Optimal emission *e*^∗^ (dark green) and response threshold *α*^∗^ (light green) as functions of leaf damage per herbivory *d*. (B) Corresponding defense probabilities under herbivory *ψ*(*e*^∗^, *α*^∗^) (brown) and in its absence *ψ*(0, *α*^∗^) (yellow). (C) Optimal emission and response threshold as functions of herbivory frequency, *ρ* (D) Corresponding defense probabilities under herbivory and in its absence. Parameters are *ρ* = 0.2, *p* = 0.5, *r* = 0.12, *c* = 0.1, and *s* = 5 in A and B, and *d* = 0.6, *p* = 0.5, *r* = 0.12, *c* = 0.1, and *s* = 5 in C and D.

### 3.3. Evolutionary condition for volatile-mediated within-plant signaling

Although solutions given by equations (20) and (21) satisfy the first-order optimality conditions, they correspond only to local optima of the loss function *L*. Therefore, the globally optimal strategy must be determined by comparing the values of *L* among all candidate strategies, including boundary solutions. Specifically, we compare the loss function *L* for signaling strategy (*e*^∗^,*α*^∗^), silent strategy (0,∞), and constitutive defense strategy (0, −∞), which are given by

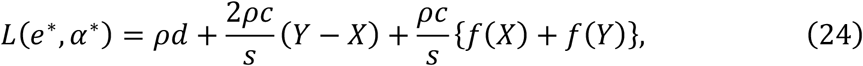

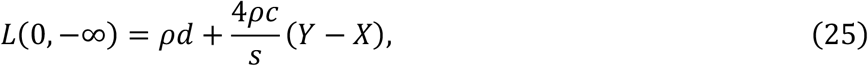

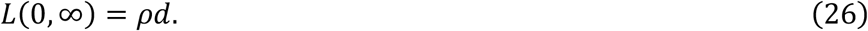

Here, *f*(*x*)is a monotonically decreasing function of *x* ≥ 1 defined as

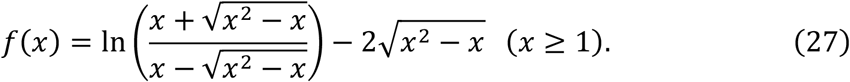

When *Y*−*X*<0, corresponding to the case of a small benefit of conditional response, the signaling strategy (*e*^∗^,*α*^∗^) and the constitutive defense strategy (0,−∞) compete for global optimality, depending on the balance between the second and third terms in equation (24). In contrast, when *Y*−*X*>0, the signaling strategy (*e*^∗^,*α*^∗^) and the silent strategy (0,∞) become the relevant candidates for the globally optimal solution. We can derive these boundaries using the approximation of *f*(*x*) in the case of |*X*-*Y*|<<1 or *X, Y* >>1 (see Appendix). The condition where the signaling strategy becomes globally optimal is summarized in Figure 4. Three boundaries separate three regions corresponding to one of the optimal strategies (Fig. 4). The boundary between regions for the silent strategy and the constitutive defense strategy is given by *X* = *Y*. In the region where damage *d* is larger than *d*_*c*_ determined by equation (16), and frequency *ρ* is smaller than *ρ*_*c*_ determined by equation (17), the signaling strategy becomes locally optimal. Within the region, the colored area represents the parameter region where the signaling strategy becomes the global optimum. The boundaries separating signaling strategy from the other two strategies are given by *L*(*e*^∗^,*α*^∗^) = *L*(0,∞) and *L*(*e*^∗^,*α*^∗^) = *L*(0,−∞). These results demonstrate that volatile-mediated signaling evolves only when damage per herbivory event is sufficiently large and herbivory frequency is intermediate. This condition is consistent with previous theoretical predictions for the evolution of inducible defense (Ito & Sakai, 2009). Notably, when damage is moderate, increasing herbivory frequency leads to a qualitative shift in the optimal strategy from silent defense to signaling, and eventually to constitutive defense.

**Figure 4.**
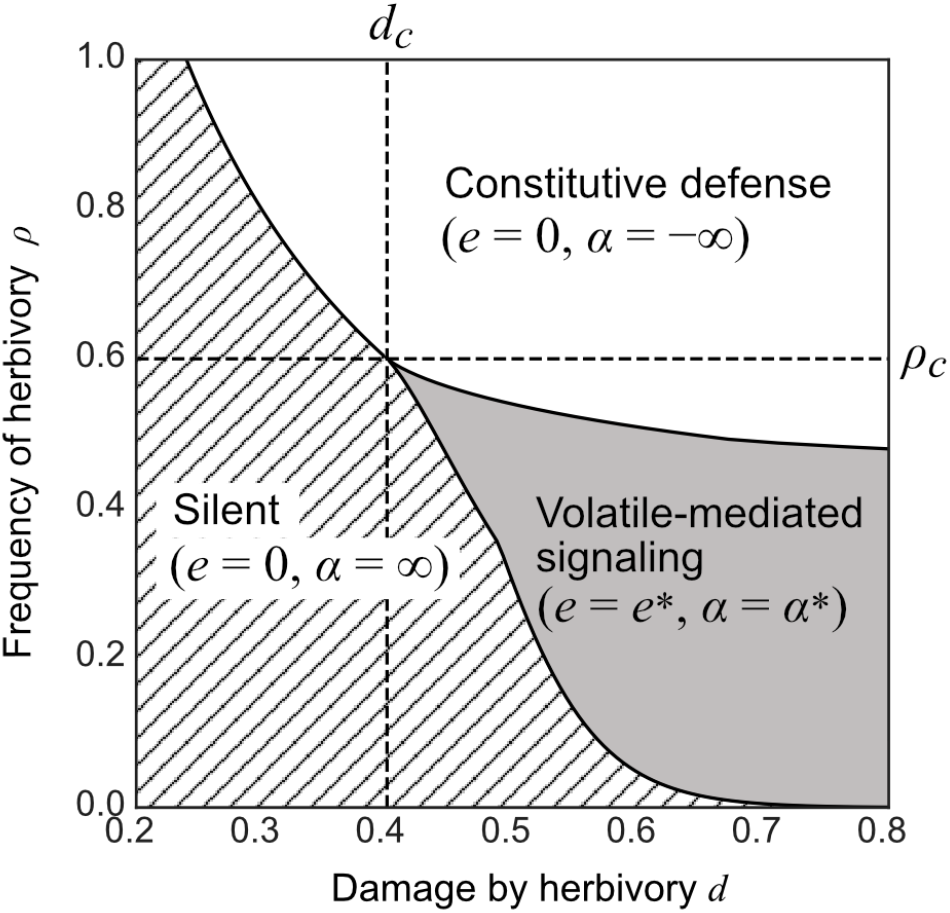
Parameter regions for three defense strategies illustrated on a *d*−*ρ* plane. The boundary between regions for the silent strategy and the constitutive defense strategy is given by *X* = *Y*. In the region where damage *d* is larger than *d*_*c*_ determined by Eqs(16), and frequency *ρ* is smaller than *ρ*_*c*_ determined by Eqs(17), the signaling strategy becomes local optimal. The boundaries defining the region where signaling strategy becomes global optimal is given by *L*(*e*^∗^,*α*^∗^) = *L*(0,∞) and *L*(*e*^∗^,*α*^∗^) = *L*(0,−∞). Parameters are *p* = 0.5, *r* = 0.12, *c* = 0.1, and *s* = 5.

Cost for signal production *c* and that for defense response *r* further influence the optimality of volatile-mediated strategies (Fig. 5). Increasing the cost of signal production *c* restricts the conditions under which signaling is favored by raising the threshold level of herbivory damage *d* required for signaling to become optimal (Fig. 5A). Under high herbivory frequency, emitting volatiles becomes inefficient, and constitutive defense is favored instead. As a result, the range of attack frequencies *ρ* over which signaling-mediated inducible defense is optimal narrows sharply with increasing signaling cost (Fig. 5B). In contrast, increasing the cost of defense *r* also raises the required damage threshold (Fig. 5C), but its effect on herbivory frequency differs. When defense costs are moderate, the range of frequencies favoring signaling-mediated inducible defense becomes widest (Fig. 5D). This occurs because higher defense costs render constitutive defense inefficient, thereby promoting inducible strategies despite their signaling costs.

**Figure 5.**
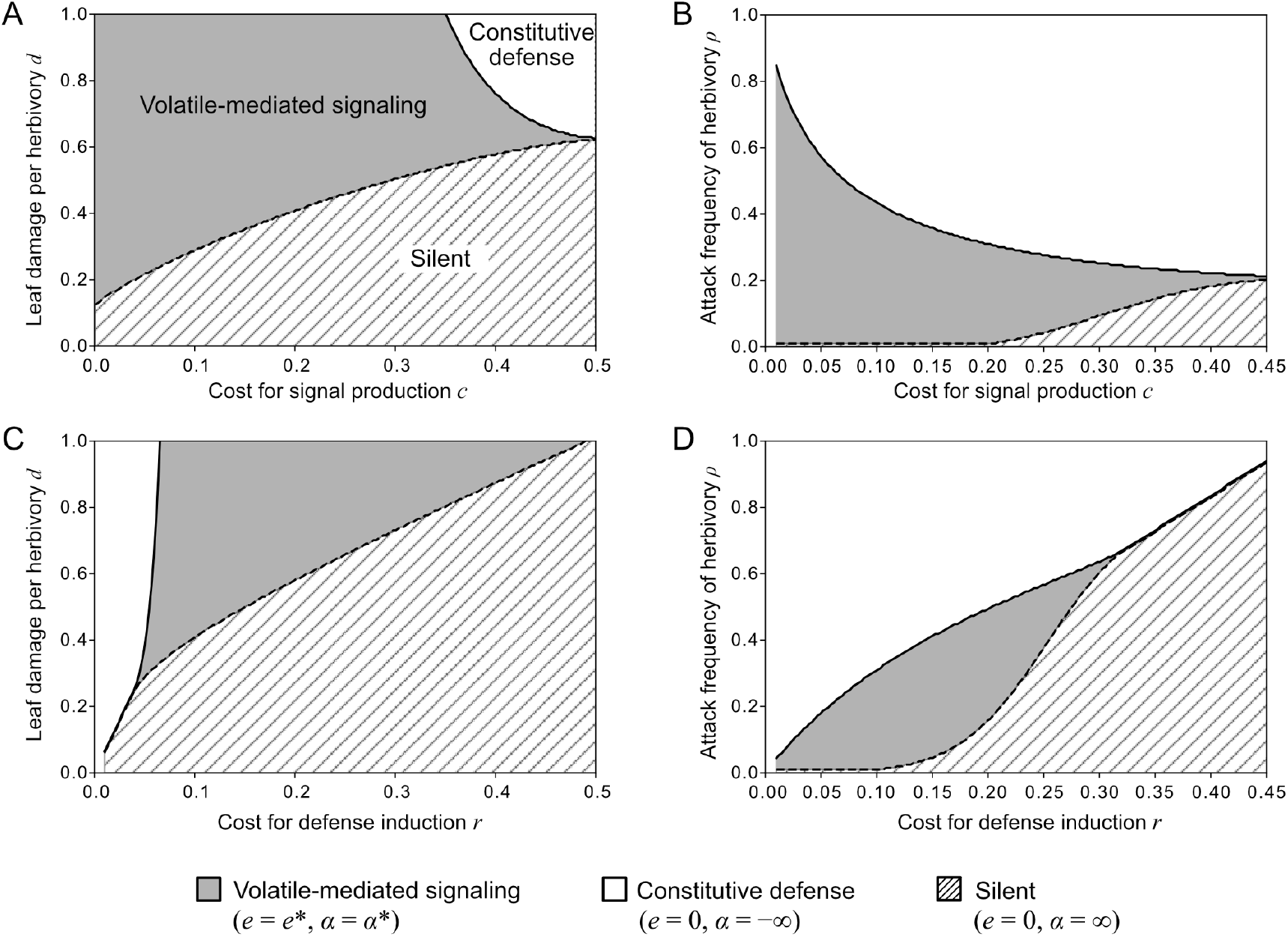
Effect of costs for signal production and defense response on the evolutionary condition of HIPV-mediated signaling. (A–B) Dependence of optimal defense strategies on signaling cost *c* (A) Parameter regions in the plane of damage per herbivory event *d* and signaling cost *c* (*ρ* = 0.2, *p* = 0.8, *r* = 0.1, *s* = 5). (B) Parameter regions in the plane of herbivory frequency *ρ* and signaling cost *c* (*d* = 0.6, *p* = 0.8, *r* = 0.1, *s* = 5). (C–D) Dependence of optimal strategies on defense induction cost *r* (C) Parameter regions in the *d*–*r* plane (*ρ* = 0.2, *p* = 0.8, *c* = 0.2, *s* = 5). (D) Parameter regions in the *ρ*−*r* plane (*d* = 0.6, *p* = 0.8, c= 0.2, *s* = 5).

### 3.4. Effects of perception-independent herbivory suppression by volatile emission

In the previous sections, we assumed that volatile emission contributes to plant fitness exclusively through the induction of defense responses. However, plant volatiles are also known to reduce herbivory independently of systemic defense induction, for example by repelling herbivores (De Moraes et al., 2001) or attracting natural enemies (Takabayashi, 2022; Takabayashi & Dicke, 1996). Such dual defensive functions of plant volatiles may promote the evolution of volatile-mediated signaling and help explain the widespread occurrence of volatile emission across plant taxa (Kessler, 2015). To examine how such a perception-independent defensive effect influences the evolution of volatile-mediated signaling strategies, we extended the original model by incorporating a mechanism in which herbivory damage is reduced by a fixed proportion depending on the amount of volatile emission as follows:

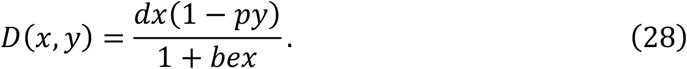

Hereafter, we refer to this extended model as the bifunctional model. Figure 6 compares the original signaling model with the bifunctional model. The bifunctional model predicts that volatile-emitting strategies become optimal over a substantially broader range of herbivory damage, herbivory frequency, and signaling costs (Fig. 6). In addition, the bifunctional model generates new strategies among volatile-emitting plants. Besides signaling strategies that induce defense responses, two emission-only strategies emerge: volatile emission without defense induction (*e,α*) = (*e*^∗^,∞) (Fig. 6A−D) and volatile emission combined with constitutive defense (*e,α*) = (*e*^∗^,−∞) (Fig. 6B).

**Figure 6.**
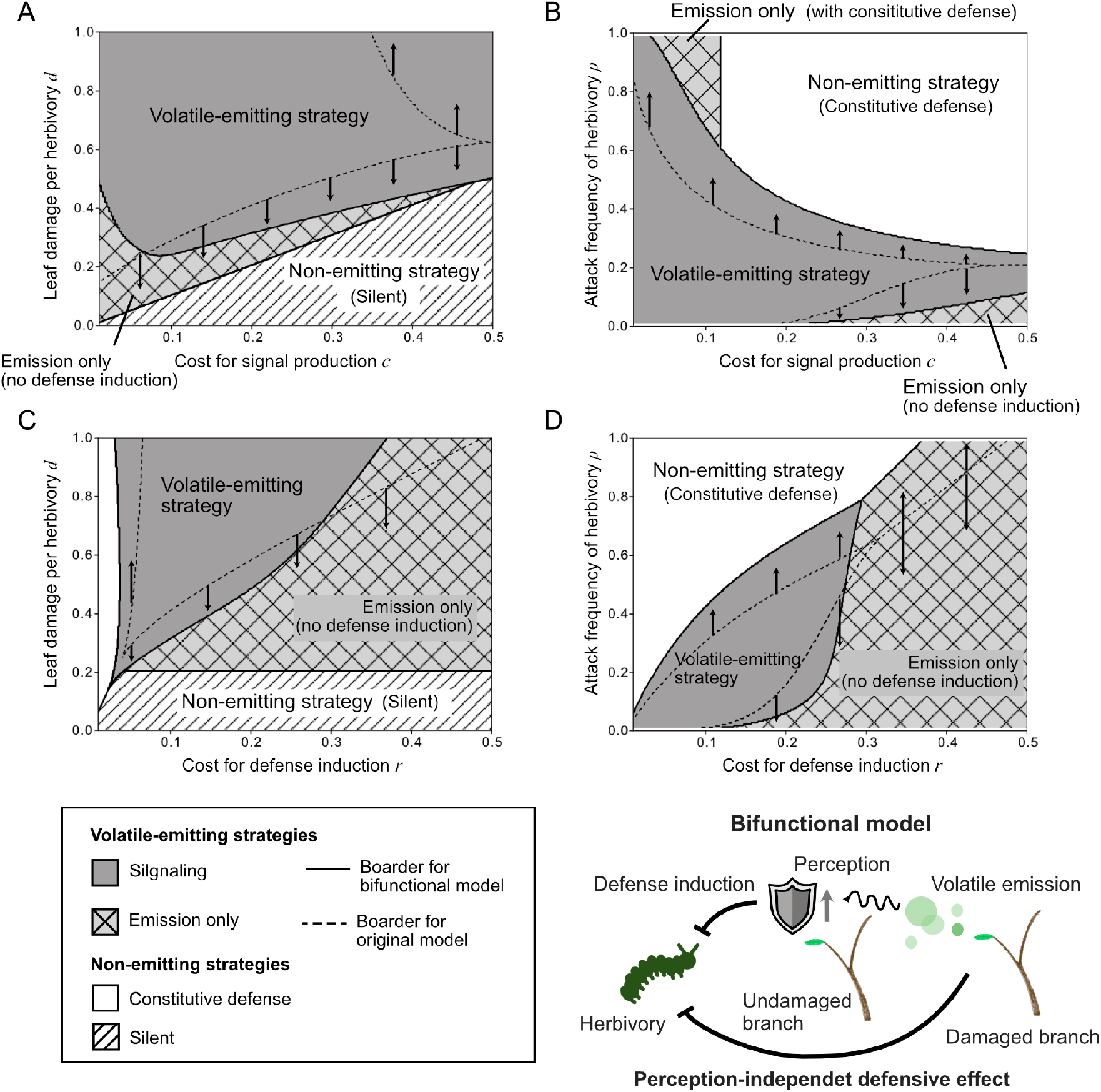
Effects of bifunctionality on the evolution of volatile emission strategies. (A– B) Dependence of optimal defense strategies on signaling cost *c*, in the *d*−*c* plane (A) and the *ρ*−*c* plane (B). (C–D) Dependence of optimal strategies on defense response cost *r*, in the *d*–*r* plane (C) and the *ρ*−*r* plane (D). Dark gray and light gray regions represent HIPV-emitting strategies. In particular, light gray region with cross-hatching indicates the emission-only strategy. In addition, white and diagonal hatched (//) region represent constitutive defense and silent strategies, respectively. The boundary between silent and emission-only strategies is determined by Eqs (30). Relative to the original model (dashed line), the bifunctional model (solid line) expands the parameter space in which volatile emission evolves and gives rise to an emission-only strategy. Parameters are the same as those in Fig. 5, except for bifunctional model-specific parameter *b*=1.

To understand these results mathematically, we compared the loss function of the bifunctional model, *L*_)_(*e, α*)with that of the original model *L*(*e, α*).

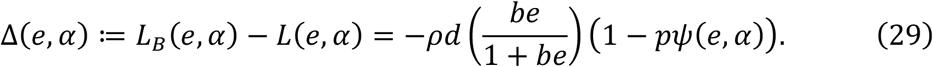

The equation (24) indicates the reduction in loss caused by bifunctionality depends on the volatile emission level *e* and the probability of defense induction *ψ*(*e, α*). Because strategies without volatile emission (silent and constitutive defense strategies) have *e* = 0, their losses remain identical in both models (*Δ*(*e, α*) = 0). In contrast, volatile-emitting strategies experience a reduction (*Δ*(*e, α*) < 0) in carbon loss in the bifunctional model. As a consequence, parameter regions in which silent or constitutive strategies were optimal in the original model shift toward volatile-emitting strategies in the bifunctional model. Furthermore, because the reduction in loss is greatest when defense induction does not occur (*α*→∞), emission-only strategies tend to emerge near the boundary between silent and signaling regions. In particular, boundary between silent *L*(0, ∞) and emission-only strategy *L*(*e*, ∞) can be derived by the comparison of their loss function:

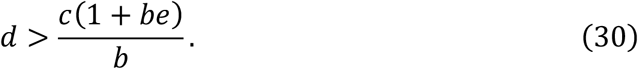

Overall, bifunctionality increases the adaptive value of volatile emission, broadens the conditions under which volatile-emitting strategies are favored, and enables the evolution of emission-only strategies, thereby decoupling volatile emission from perception.

## 4. Discussion

Our analysis of mathematical model describing the coevolutionary dynamics between emission and perception traits derived the condition under which plants exhibit both a positive volatile emission rate and a perception-dependent response to volatiles emitted by different branches (Eqs. 16 and 17). We demonstrated that increasing herbivory frequency leads to a reduction in HIPV emission and a concomitant increase in HIPV sensitivity for defense induction (Fig. 3). Furthermore, we showed that the perception-independent defensive effects of HIPVs can decouple the joint evolution of emission and perception, resulting in the emergence of a strategy that only emits HIPVs without defense induction (Fig. 5). In the following sections, we discuss these three points by comparing with previous theoretical study and empirical knowledges.

### 4.1. Three evolutionary outcomes of HIPV emission and perception traits

We showed that herbivory-associated parameters – herbivory damage *d* and frequency *ρ* – govern the optimality of HIPV-mediated signaling strategies (Fig. 3; Fig. 4). This result is qualitatively consistent with previous theoretical work demonstrating that signal-dependent inducible defenses evolve only when herbivory damage is sufficiently large and occurs at intermediate frequencies (Ito & Sakai, 2009). Building on this finding, we further demonstrated that this evolutionary condition can be succinctly characterized by two quantities: the relative benefit of defense *X* and the relative benefit of inducibility *Y* (Eqs. 16 and 17). The relative benefit of defense is defined as the net gain from defense, given by the reduction in damage *pd* minus the defense cost *r*. In contrast, the relative benefit of inducibility is defined as the cost of unnecessary defense in the absence of herbivory, *r*(1 - *ρ*), normalized by herbivory frequency *ρ*. This quantity declines as herbivory becomes more frequent, reflecting the reduced duration of herbivore-free periods. Based on these measures, the coevolutionary outcomes of emission and perception traits can be summarized as follows. A signaling strategy is favored only when both relative benefits are sufficiently large. When herbivory damage is small, the relative benefit of defense is low, and a silent strategy—characterized by the absence of defense— is optimal. Conversely, as herbivory frequency increases, the relative benefit of inducibility declines, favoring a constitutive defense strategy that does not rely on HIPV signaling.

Empirical examples corresponding to the silent strategy, characterized by reduced investment in defense, are supported by studies demonstrating trade-offs between growth and defense. Reduced herbivore pressure, for instance through insect-removal experiments, can relax herbivore-mediated selection, leading to decreased levels of defensive compounds and increased competitive ability (Agrawal et al., 2012). In contrast, constitutive defense is generally defined as defense expressed irrespective of herbivore attack and includes both physical barriers—such as surface waxes, the cuticle, and trichomes—and chemical defenses, including various secondary metabolites such as alkaloids and glucosinolates (Agrawal, 2007; Gatehouse, 2002). A representative example of systems dominated by constitutive defense is found in tropical plant species. Consistent with our theoretical predictions, defensive secondary metabolite production in *Inga* species from Neotropical rainforests—where herbivore pressure is persistently high—has been shown to be largely unresponsive to herbivory (Bixenmann et al., 2016). However, it is important to note that these studies do not explicitly test whether plants employ HIPV-mediated within-plant signaling. In sagebrush, *Artemisia cana*, a close relative of Artemisia tridentata—a well-known species that uses HIPV-mediated within-plant signaling (Karban et al., 2006)—exhibits systemic defense responses that do not rely on volatile signaling (Shiojiri & Karban, 2008). Such inter-specific variation provides a valuable opportunity to link differences in dependence on HIPV-mediated within-plant signaling with the evolution of defense strategies.

### 4.2. Joint evolution of reduced HIPV emission and enhanced sensitivity to HIPVs

Within the parameter regime in which the signaling strategy is favored, our results indicate that increasing herbivory frequency leads to reduced HIPV emission and heightened sensitivity to HIPVs (Fig. 3). This pattern is advantageous because it can reduce the cost for HIPV signaling per herbivory event while maintaining a high inducibility of defense response. However, a lower threshold for HIPV response also increase the likelihood of unnecessary defense induction in the absence of herbivory. Therefore, this signaling strategy is selected only when herbivory frequency is moderate (Fig. 4). This result may help explain empirical findings from meta-analyses showing that cultivated plants—expected to experience weaker herbivore-driven selection—often exhibit higher levels of HIPV emission than their wild plants (Rowen & Kaplan, 2016), although this study compared HIPV emission only, and did not directly assess its role in within-plant signaling.

An important empirical study to compare our findings is provided by field manipulation experiments involving artificial insect removal in *Solidago altissima* populations (Kalske et al., 2019). In this study, individuals from herbivore-exposed populations responded broadly to volatiles emitted by damaged plants, whereas those from herbivore-free populations responded only to volatiles from genetically identical individuals. Although this study focuses on plant–plant communication, such broadened responsiveness is consistent with a lowered response threshold for HIPVs, as predicted by our model, which would increase responsiveness not only to within-plant cues but also to volatiles from neighboring individuals. In contrast, the study suggested that the evolution of sensitivity was not uniform; rather, it varied with genetic relatedness, indicating a role for kin selection in shaping HIPV-mediated communication (Kalske et al., 2019). Because our model considers signaling among branches within a single individual, extending the framework to incorporate interactions among individuals with varying degrees of relatedness would provide a valuable avenue for investigating how social context influences the coevolution of HIPV emission and perception traits. With respect to HIPV emission, no significant difference in total emission was observed between treatments (Kalske et al., 2019), which is not fully consistent with our model predictions. Instead, herbivore presence reduced variation in HIPV composition within populations, indicating the evolution of information-sharing signals. Such compositional shifts cannot be captured by our current model, which treats HIPV emission as a single quantitative trait. These discrepancies between empirical and theoretical findings highlight the need for an extended modeling framework that incorporates a “blend” of multiple chemical components to describe HIPV-mediated signaling system.

### 4.3. Decoupling the joint evolution of emission and perception by perception-independent defensive effects of HIPVs

Our model shows that defensive benefits obtained solely from HIPV emission—such as predator attraction or herbivore deterrence—expand the parameter space in which HIPV-mediated signaling can evolve, while also allowing conditions under which emission alone becomes the optimal strategy (Fig. 6). This result predicts the existence of plants that emit HIPVs but exhibit relatively weak responsiveness to them. However, there is currently no clear empirical evidence for the existence of such phenotypes. This is partly because, despite the extensive body of research on plant–insect and plant–plant communication, studies that directly demonstrate within-plant signaling remain relatively limited (Li & Blande, 2017), even though these forms of communication are thought to have derived from intra-plant signaling processes (Heil & Karban, 2010). To test our theoretical predictions, future studies should comprehensively quantify not only HIPV emission but also responsiveness to HIPVs released by the same individuals.

In addition, while our model assumes the case that benefits are biased toward HIPV emission, it is also possible for plants to gain benefits solely through perception. One such scenario is plant–plant communication, in which individuals “eavesdrop” on HIPVs emitted by neighboring plants without producing volatiles themselves. Indeed, a theoretical study on plant-plant communication based on evolutionary game theory suggested that when the benefits of inter-plant signaling exceed those of within-plant signaling, non-emitting “cheater” strategies can evolve (Hirose & Satake, 2024). Such perception-biased benefits may counterbalance the emission-biased benefits considered in this study and thereby contribute to the maintenance of HIPV-mediated signaling under certain ecological conditions. To evaluate this possibility, our coevolutionary model of HIPV emission and perception should incorporate volatile cues derived from neighboring individuals, thereby situating the framework within the context of plant–plant communication.

### 4.4. Conclusion and future perspective

In this study, we theoretically analyzed the joint evolution of HIPV emission and perception in the context of within-plant signaling. Although our model was developed for within-plant communication, its framework is not restricted to this level and will provide a general basis for understanding the evolution of HIPV-mediated interaction such as plant–plant communication. In addition, recent advances in molecular mechanism of HIPV emission and perception may facilitate the development of more precise and predictive models. In particular, metabolic pathways and genes involved in the biosynthesis of plant volatiles have been increasingly characterized (Arimura et al., 2009), enabling comparative genomic approaches to investigate the evolutionary gain and loss of emission capacity across species (Ikezaki et al., 2025; Shang et al., 2024). In contrast, the molecular mechanisms of HIPV perception remain less well understood; however, recent studies have identified several transcriptional regulators (Nagashima et al., 2019) and signal transduction mechanisms in response to volatiles (Aratani et al., 2023). Incorporating such molecular insights into mathematical models would introduce physiologically and metabolically grounded constraints on the evolution of HIPV emission and perception. This integration would deepen our understanding of both the proximate and ultimate drivers of the evolution of HIPV-mediated signaling.

## 5. Acknowledgements

We would like to thank T Hagiwara and S Hirose for valuable comments and discussion.

This work was supported by JST SPRING, Japan Grant Number JPMJSP2136.

## 6. Author Contribution

**Shuichi N Kudo**: Writing – original draft, Writing – review and editing, Conceptualization, Formal analysis, Investigation, Methodology, Validation, Project administration, Funding acquisition. **Koki Iwakura**: Writing – original draft, Writing – review and editing, Conceptualization, Formal analysis, Investigation, Methodology, Validation. **Akiko Satake**: Writing – review and editing, Conceptualization, Project administration, Supervision.

## 9.Appendix

### Condition for global optimality of the volatile-mediated signaling strategy

When Y−X<0 (that is, the benefit of conditional response is small), the signaling strategy (*e*^∗^,*α*^∗^) and the constitutive defense strategy (0,−∞) compete for global optimality, depending on the balance between the second and third terms in (19). Comparing *L*(*e*^*\**^, *α*^*^) and *L*(0, -∞), we obtain a condition under which the signaling strategy is globally optimal. Define

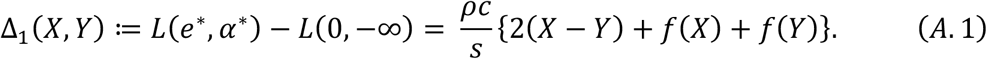

Solving Δ_%_(*X,Y*) ≤0 yields

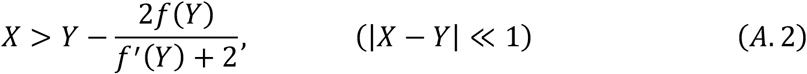

and

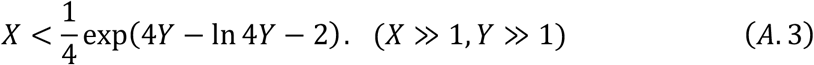

To obtain these inequalities, we use the approximation

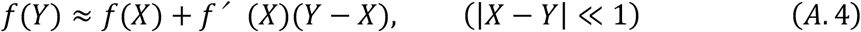

and

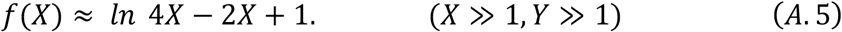

The first approximation follows from the Taylor expansion of *f* about *X* around *Y*, while the second is obtained from the Taylor expansion of *f* after changes of variables. For the case that |*X* −*Y*| ≪1, substituting (A. 4) into (A. 1) gives

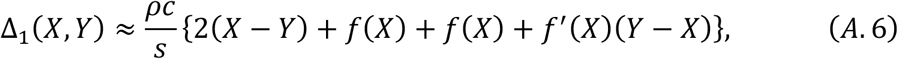

which yields (A. 2). For the case that *X* ≫ 1 and *Y* ≫ 1, using (A. 5) gives

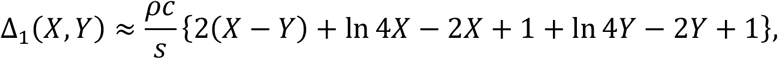

which yields (A. 3).

In contrast, when *Y* – *X>0*, the candidates of the globally optimal strategy are the signaling strategy (*e*\*, *α*^*^) and the silent strategy (0, ∞). Comparing *L*(*e*\*, *α*\*) and *L*(0, ∞), we obtain the condition where the signaling strategy becomes globally optimal. Define

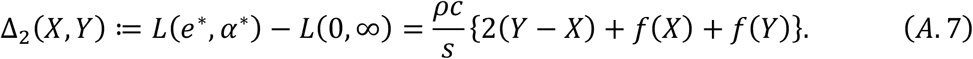

By solving Δ_1_(*X,Y*)≤0yields

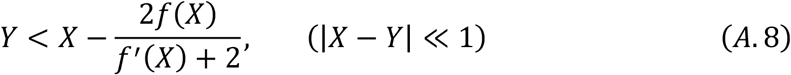

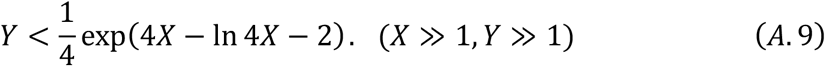

These inequalities are derived using the same approximations as above.

## References

Agrawal, A. A. (2007). Macroevolution of plant defense strategies. in Trends in Ecology and Evolution (Vol. 22, Number 2, pp. 103–109). 10.1016/j.tree.2006.10.012

Agrawal, A. A., Hastings, A. P., Johnson, M. T. J., Maron, J. L., & Salminen, J.-P. (2012). Insect Herbivores Drive Real-Time Ecological and Evolutionary Change in Plant Populations. Science, 338(6103), 113–116. 10.1126/science.1225977

Aratani, Y., Uemura, T., Hagihara, T., Matsui, K., & Toyota, M. (2023). Green leaf volatile sensory calcium transduction in Arabidopsis. Nature Communications, 14(1). 10.1038/s41467-023-41589-9

Arimura, G. I., Matsui, K., & Takabayashi, J. (2009). Chemical and molecular ecology of herbivore-induced plant volatiles: Proximate factors and their ultimate functions. in Plant and Cell Physiology (Vol. 50, Number 5, pp. 911–923). 10.1093/pcp/pcp030

Bixenmann, R. J., Coley, P. D., Weinhold, A., & Kursar, T. A. (2016). High herbivore pressure favors constitutive over induced defense. Ecology and Evolution, 6(17), 6037–6049. 10.1002/ece3.2208

Blossey, B., & Notzold, R. (1995). Evolution of Increased Competitive Ability in Invasive Nonindigenous Plants: A Hypothesis. in Source: Journal of Ecology (Vol. 83, Number 5).

Boake, C. R. B. (1991). Coevolution of senders and receivers of sexual signals: Genetic coupling and genetic correlations. Trends in Ecology & Evolution, 6(7), 225–227. 10.1016/0169-5347(91)90027-U

Charlton, B. D., Owen, M. A., & Swaisgood, R. R. (2019). Coevolution of vocal signal characteristics and hearing sensitivity in forest mammals. Nature Communications, 10(1). 10.1038/s41467-019-10768-y

De Moraes, C. M., Mescher, M. C., & Tumlinson, J. H. (2001). Caterpillar-induced nocturnal plant volatiles repel conspeci®c females. Nature, 410. 10.1038/35069058

Dicke, M., & Baldwin, I. T. (2010). The evolutionary context for herbivore-induced plant volatiles: beyond the “cry for help.” In Trends in Plant Science (Vol. 15, Number 3, pp. 167–175). 10.1016/j.tplants.2009.12.002

Dudareva, N., Pichersky, E., & Gershenzon, J. (2004). Biochemistry of plant volatiles. in Plant Physiology (Vol. 135, Number 4, pp. 1893–1902). American Society of Plant Biologists. 10.1104/pp.104.049981

Fagerström, T., Larsson, S., & Tenow, O. (1987). On Optimal Defence in Plants. in Source: Functional Ecology (Vol. 1, Number 2).

Frost, C. J., Appel, H. M., Carlson, J. E., De Moraes, C. M., Mescher, M. C., & Schultz, J. C. (2007). Within-plant signalling via volatiles overcomes vascular constraints on systemic signalling and primes responses against herbivores. Ecology Letters, 10(6), 490–498. 10.1111/j.1461-0248.2007.01043.x

Frost, C. J., Mescher, M. C., Carlson, J. E., & De Moraes, C. M. (2008). Plant defense priming against herbivores: Getting ready for a different battle. in Plant Physiology (Vol. 146, Number 3, pp. 818–824). American Society of Plant Biologists. 10.1104/pp.107.113027

Gatehouse, J. A. (2002). Plant resistance towards insect herbivores: A dynamic interaction. in New Phytologist (Vol. 156, Number 2, pp. 145–169). 10.1046/j.1469-8137.2002.00519.x

Gouinguené, S., Degen, T., & Turlings, T. C. J. (2001). Variability in herbivore-induced odour emissions among maize cultivars and their wild ancestors (teosinte). Chemoecology, 11(1), 9–16. 10.1007/PL00001832

Groot, A. T., Dekker, T., & Heckel, D. G. (2016). The Genetic Basis of Pheromone Evolution in Moths. in Annual Review of Entomology (Vol. 61, pp. 99–117). Annual Reviews Inc. 10.1146/annurev-ento-010715-023638

Hare, J. D. (2007). Variation in herbivore and methyl jasmonate-induced volatiles among genetic lines of Datura wrightii. Journal of Chemical Ecology, 33(11), 2028–2043. 10.1007/s10886-007-9375-1

Heil, M., & Karban, R. (2010). Explaining evolution of plant communication by airborne signals. Trends in Ecology and Evolution, 25(3), 137–144. 10.1016/j.tree.2009.09.010

Heil, M., & Silva Bueno, J. C. (2007). Within-plant signaling by volatiles leads to induction and priming of an indirect plant defense in nature. in PNAS (Vol. 104). 10.1073pnas.0610266104

Hirose, S., & Satake, A. (2024). Theoretical analyses for the evolution of biogenic volatile organic compounds (BVOC) emission strategy. Ecology and Evolution, 14(7). 10.1002/ece3.11548

Huang, W., Siemann, E., Wheeler, G. S., Zou, J., Carrillo, J., & Ding, J. (2010). Resource allocation to defence and growth are driven by different responses to generalist and specialist herbivory in an invasive plant. Journal of Ecology, 98(5), 1157–1167. 10.1111/j.1365-2745.2010.01704.x

Ikezaki, Y., Kudo, S. N., Nakata, T., Koita, S., Munakata, R., Yazaki, K., Torimaru, T., Tomaru, N., Isobe, S., Hirakawa, H., Kusumi, J., & Satake, A. (2025). Molecular evolution of terpene synthase underlying the diversification of isoprene emission in Fagaceae. 10.1101/2025.07.31.667835

Ito, K., & Sakai, S. (2009). Optimal defense strategy against herbivory in plants: Conditions selecting for induced defense, constitutive defense, and no-defense. Journal of Theoretical Biology, 260(3), 453–459. 10.1016/j.jtbi.2009.07.002

Iwasa, Y., Hayashi, R., & Satake, A. (2025). Optimal seasonal schedule for producing biogenic volatile organic compounds for tree defense. Journal of Theoretical Biology, 596. 10.1016/j.jtbi.2024.111986

Kalske, A., Shiojiri, K., Uesugi, A., Sakata, Y., Morrell, K., & Kessler, A. (2019). Insect Herbivory Selects for Volatile-Mediated Plant-Plant Communication. Current Biology, 29(18), 3128-3133.e3. 10.1016/j.cub.2019.08.011

Karban, R., Shiojiri, K., Huntzinger, M., & Mccall, A. C. (2006). DAMAGE-INDUCED RESISTANCE IN SAGEBRUSH: VOLATILES ARE KEY TO INTRA-AND INTERPLANT COMMUNICATION. in Ecology (Vol. 87, Number 4).

Kessler, A. (2015). The information landscape of plant constitutive and induced secondary metabolite production. in Current Opinion in Insect Science (Vol. 8, pp. 47–53). Elsevier Inc. 10.1016/j.cois.2015.02.002

Kobayashi, Y., & Yamamura, N. (2003). Evolution of signal emission by non-infested plants growing near infested plants to avoid future risk. Journal of Theoretical Biology, 223(4), 489–503. 10.1016/S0022-5193(03)00124-3

Kobayashi, Y., Yamamura, N., & Sabelis, M. W. (2006). Evolution of talking plants in a tritrophic context: Conditions for uninfested plants to attract predators prior to herbivore attack. Journal of Theoretical Biology, 243(3), 361–374. 10.1016/j.jtbi.2006.05.026

Li, T., & Blande, J. D. (2017). How common is within-plant signaling via volatiles? Plant Signaling and Behavior, 12(8). 10.1080/15592324.2017.1347743

Lin, T., Vrieling, K., Laplanche, D., Klinkhamer, P. G. L., Lou, Y., Bekooy, L., Degen, T., Bustos-Segura, C., Turlings, T. C. J., & Desurmont, G. A. (2021). Evolutionary changes in an invasive plant support the defensive role of plant volatiles. Current Biology, 31(15), 3450-3456.e5. 10.1016/j.cub.2021.05.055

Loughrin, J. H., Manukian, A., Heath, R. R., & Tumlinson, J. H. (1995). VOLATILES EMITTED BY DIFFERENT COTTON VARIETIES DAMAGED BY FEEDING BEET ARMYWORM LARVAE. in Journal of Chemical Ecology (Vol. 21, Number 8).

McKey, D. (1974). Adaptive Patterns in Alkaloid Physiology. The American Naturalist, 108, 305–320.

Nagashima, A., Higaki, T., Koeduka, T., Ishigami, K., Hosokawa, S., Watanabe, H., Matsui, K., Hasezawa, S., & Touhara, K. (2019). Transcriptional regulators involved in responses to volatile organic compounds in plants. Journal of Biological Chemistry, 294(7), 2256–2266. 10.1074/jbc.RA118.005843

Niinemets, Üi. (2018). Storage of defense metabolites in the leaves of Myrtaceae: News of the eggs in different baskets. in Tree Physiology (Vol. 38, Number 10, pp. 1445– 1450). Oxford University Press. 10.1093/treephys/tpy115

Rowen, E., & Kaplan, I. (2016). Eco-evolutionary factors drive induced plant volatiles: A meta-analysis. New Phytologist, 210(1), 284–294. 10.1111/nph.13804

Schuman, M. C., Heinzel, N., Gaquerel, E., Svatos, A., & Baldwin, I. T. (2009). Polymorphism in jasmonate signaling partially accounts for the variety of volatiles produced by Nicotiana attenuata plants in a native population. New Phytologist, 183(4), 1134–1148. 10.1111/j.1469-8137.2009.02894.x

Shang, J., Feng, D., Liu, H., Niu, L., Li, R., Li, Y., Chen, M., Li, A., Liu, Z., He, Y., Gao, X., Jian, H., Wang, C., Tang, K., Bao, M., Wang, J., Yang, S., Yan, H., & Ning, G. (2024). Evolution of the biosynthetic pathways of terpene scent compounds in roses. Current Biology, 34(15), 3550-3563.e8. 10.1016/j.cub.2024.06.075

Sharkey, T. D., Wiberley, A. E., & Donohue, A. R. (2008). Isoprene emission from plants: Why and how. in Annals of Botany (Vol. 101, Number 1, pp. 5–18). 10.1093/aob/mcm240

Shiojiri, K., & Karban, R. (2008). Vascular Systemic Induced Resistance for Artemisia cana and Volatile Communication for Artemisia douglasiana. in Source: The American Midland Naturalist (Vol. 159, Number 2).

Strauss, S. Y., Rudgers, J. A., Lau, J. A., & Irwin, R. E. (2002). Direct and ecological costs of resistance to herbivory. in TRENDS in Ecology & Evolution (Vol. 17, Number 6). http://tree.trends.com0169-5347/02/$-seefrontmatter

Symonds, M. R. E., & Elgar, M. A. (2008). The evolution of pheromone diversity. in Trends in Ecology and Evolution (Vol. 23, Number 4, pp. 220–228). 10.1016/j.tree.2007.11.009

Takabayashi, J. (2022). Herbivory-Induced Plant Volatiles Mediate Multitrophic Relationships in Ecosystems. Plant and Cell Physiology, 63(10), 1344–1355. 10.1093/pcp/pcac107

Takabayashi, J., & Dicke, M. (1996). Plant-carnivore mutualism through herbivore-induced carnivore attractants. Trends in Plant Science, 1, 109–113. 10.1016/S1360-1385(96)90004-7

War, A. R., Sharma, H. C., Paulraj, M. G., War, M. Y., & Ignacimuthu, S. (2011). Herbivore induced plant volatiles: Their role in plant defense for pest management. Plant Signaling & Behavior, 6(12), 1973–1978. 10.4161/psb.6.12.18053

Yamauchi, A., Van Baalen, M., Kobayashi, Y., Takabayashi, J., Shiojiri, K., & Sabelis, M. W. (2015). Cry-wolf signals emerging from coevolutionary feedbacks in a tritrophic system. Proceedings of the Royal Society B: Biological Sciences, 282(1818). 10.1098/rspb.2015.2169

